# LYRUS: A Machine Learning Model for Predicting the Pathogenicity of Missense Variants

**DOI:** 10.1101/2021.05.10.443497

**Authors:** Jiaying Lai, Jordan Yang, Ece D. Gamsiz Uzun, Brenda M. Rubenstein, Indra Neil Sarkar

## Abstract

Single amino acid variations (SAVs) are a primary contributor to variations in the human genome. Identifying pathogenic SAVs can aid in the diagnosis and understanding of the genetic architecture of complex diseases, such as cancer. Most approaches for predicting the functional effects or pathogenicity of SAVs rely on either sequence or structural information. Nevertheless, previous analyses have shown that methods that depend on only sequence or structural information may have limited accuracy. Recently, researchers have attempted to increase the accuracy of their predictions by incorporating protein dynamics into pathogenicity predictions. This study presents < Lai Yang Rubenstein Uzun Sarkar > (LYRUS), a machine learning method that uses an XGBoost classifier selected by TPOT to predict the pathogenicity of SAVs. LYRUS incorporates five sequence-based features, six structure-based features, and four dynamics-based features. Uniquely, LYRUS includes a newly-proposed sequence co-evolution feature called variation number. LYRUS’s performance was evaluated using a dataset that contains 4,363 protein structures corresponding to 20,307 SAVs based on human genetic variant data from the ClinVar database. Based on our dataset, the LYRUS classifier has a higher accuracy, specificity, F-measure, and Matthews correlation coefficient (MCC) than alternative methods including PolyPhen2, PROVEAN, SIFT, Rhapsody, EVMutation, MutationAssessor, SuSPect, FATHMM, and MVP. Variation numbers used within LYRUS differ greatly between pathogenic and neutral SAVs, and have a high feature weight in the XGBoost classifier employed by this method. Applications of the method to PTEN and TP53 further corroborate LYRUS’s strong performance. LYRUS is freely available and the source code can be found at https://github.com/jiaying2508/LYRUS.

## Introduction

Recent technological advances such as high-throughput screening methods have made an abundance of sequencing data that have transformed our understanding of human genetic variation readily available. Since the determination of the first human genome sequence, more than one million human genomes have been collectively sequenced across the academic, clinical, and private sectors^1,2^. This increase in genomic data is revealing a growing number of rare variants, for which there is insufficient data to decipher whether they are pathogenic. Rationalizing the functional and clinical implications of these millions of observed sequence variants remains a formidable undertaking.

In the postgenomic era, understanding the relationship among genetic and phenotypic variations represents a major challenge^3^. Single nucleotide polymorphisms (SNP) refer to a single nucleotide substitution that occurs in more than 1% of the population. SNPs, which occur approximately once in every 1000 nucleotides, significantly contribute to human genetic variation and diversity^4^. There are approximately 4 to 5 million SNPs in the human genome, which result in a wide range of phenotypic properties, such as eye color, disease, and individual drug responses^5^. There are two types of coding region SNPs: (1) synonymous and (2) non-synonymous. Synonymous SNPs do not alter the encoded protein sequence, yet perturb splicing, regulatory mechanisms, and gene expression levels ^6^. Non-synonymous SNPs, on the other hand, alter the protein sequence, and may result in single amino acid variants (SAVs)^7^. Among the known disease variants, roughly 45% are missense variants that encode a single amino acid change in the affected protein^8^, which are tied to human diseases such as Parkinson’s disease, Alzheimer’s disease, and cancer^9,10^. Differentiating pathogenic SAVs from neutral SAVs is thus of great importance in the post-genomic era, as it can enhance our understanding of the correlation between genotype and phenotype, facilitating the development of novel treatment strategies for complex diseases.

The accurate classification of effects of genetic variants on various disorders remains a difficult goal to achieve, despite the abundance of genomic data collected over the last decade and the multiple efforts to elucidate their links to phenotypic traits. Most existing software for predicting the functional effects of amino acid variations are based on the assumption that protein sequences observed among living organisms have survived natural selection. As a result, evolutionarily-conserved amino acid positions across multiple species are assumed to be functionally important, and amino acid variations observed at conserved positions are assumed to be pathogenic^11^.

Previous analyses have shown that methods incorporating only sequence-related information may suffer from reduced accuracy^12^. Furthermore, Sunyaev *et al.*^13^ have shown that pathogenic mutations often affect the intrinsic structural features of proteins, including sites involved in disulphide bonds, and Wang and Moult^14^ have demonstrated that most pathogenic mutations appear to affect protein stability. It is therefore evident that knowing the impact of mutations on protein stability is essential for clarifying the relationships among the structure, function, and dynamics of a given protein. Structure-based modeling approaches have lagged behind sequence-based approaches in evaluating the effects of SAVs, even though first-generation classifiers that can take 3D structures into account have shown considerable success ^15–17^. Additionally, most computational methods focus on reaching the highest variant classification accuracy rather than understanding the modifications that occur at the molecular scale, which might be crucial for the design of drugs or treatments.

Changes in folding free energies (ΔΔG_fold_) are the standard thermodynamic measures to probe the effects of mutations on protein stability and have already been demonstrated to characterize sequence and structural patterns among human pathogenic amino acid variants^18–20^. Several computational approaches have been developed to predict ΔΔG_fold_ as a means to link it to the pathogenicity of mutations^21–26^. Besides changes in folding free energies, solvent accessibility has been known to be associated with the pathogenicity of SAVs. SAVs located on the protein surface are more likely to be neutral, whereas those that are buried in the protein core are more likely to be pathogenic^11^. Accordingly, various approaches for predicting pathogenicity that rely on structural features are available, such as Bongo, which uses graph theoretic measures to evaluate the structural impacts of single point mutations^16,27,28^. Other studies have shown that structural information can provide results of comparable quality to those that use sequence and evolutionary information in predicting pathogenic SAVs^29–31^.

In addition to sequence conservation and protein structure, protein dynamics have also been proven to be useful for predicting SAV functional impacts. Ponzoni and Bahar^32^ evaluated a set of features generated by elastic network models (ENMs) of proteins to efficiently screen protein dynamics. Their study shows the utility of considering the equilibrium dynamics of the protein as a means of improving the predictive ability of current pathogenicity predictors. Other dynamic features, such as stiffness, effectiveness, and sensitivity, have also been shown to be important in pathogenicity prediction^33^. Tools that use dynamics-based features (e.g., Rhapsody) demonstrate that predictions are improved when dynamics-based and sequence-based features are combined^34^.

Picking the most suitable machine learning (ML) algorithm that can learn the most salient of these many possible features for prediction can be challenging. The Tree-based Pipeline Optimization Tool (TPOT) is an evolutionary algorithm-based automated machine learning (autoML) system that uses genetic programming (GP) to optimize a series of feature selectors, preprocessors, and ML models to maximize classification/regression accuracy and recommend an optimal pipeline^35^. TPOT has been shown to frequently outperform standard ML analyses given no *a priori* knowledge about the problem. We utilized TPOT to search the best ML pipeline for our dataset.

We introduce LYRUS, an ML-based approach that incorporates the essential properties of structural information, evolutionary conservation, and protein dynamics, to predict the pathogenicity of SAVs. We recently developed a sequence-evolutionary based concept, called variation number, which has been shown to vary significantly among pathogenic and neutral variants in BRCA1 and BRCA2 SNPs^36^. The inclusion of variation number distinguishes LYRUS from tools currently used in the field. LYRUS was trained and evaluated on a large set of human protein variations obtained from a publicly accessible database. The results suggest that sequence-based features have higher weights than structural and dynamic features. We compared our approach to PolyPhen2, PROVEAN, SIFT, Rhapsody, EVMutation, MutationAssessor, SuSPect, FATHMM, and MVP, which is a recently developed method that uses deep residual networks and has shown high prediction accuracy^15,34,37–42^. We illustrate the utility of LYRUS by applying it to phosphatase and tensin homolog (PTEN) and tumor protein 53 (TP53).

## Methods

### Training dataset

The training dataset for the ML pipeline was generated using ClinVar, which is a public archive of human variations and phenotypes^50^. Each entry in ClinVar is associated with a review score: the larger the number of review stars an entry receives up to a maximum of four, the more verified that entry has been. All of the SAVs in ClinVar with at least one review star were obtained. The SAVs in the resulting dataset were further categorized as having a pathogenicity of benign, benign/likely benign, likely benign, likely pathogenic, pathogenic/likely pathogenic, or pathogenic. Benign, benign/likely benign, and likely benign SAVs were assigned a pathogenicity score of 0, while all other SAVs were assigned a score of 1. After data cleaning, the values of each feature for each SAV were calculated (Figure 1).

**Figure 1.**
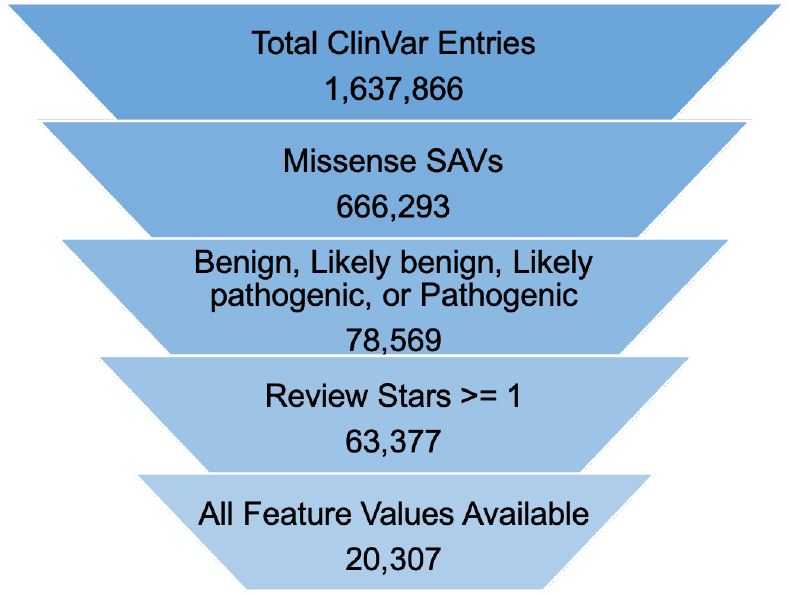
Number of SAVs from ClinVar. Among all of the SAVs available in ClinVar, roughly 1.2% of SAVs meet the selection criteria.

### Feature selection for SAV pathogenicity prediction

The three categories of features widely used in SAV pathogenicity prediction are sequence-based features, structure-based features, and dynamics-based features. We picked 15 features in total from these three categories in our prediction pipeline. The 15 features are described in Table 1 and the links to the methods used to produce them are listed in Supplementary Table S1. The variation number is a phylogenetic measure recently developed by our lab that can quantify sequence conservation using sequence homologs from different species ^36^. The pipeline for calculating variation numbers is depicted in Figure 2. The orthologous sequences required by variation number and EVMutation were obtained from the NCBI Orthologs Database. In addition to the sequence and variant information, all of the structural and dynamic features also require protein structure files from the Protein Data Bank (PDB). The PDB files were downloaded from SWISS-MODEL^51^.

**Table 1.** Features Used for SAV Pathogenicity Prediction. Fifteen features belonging to three different categories are used. Each feature calculation requires either an amino acid sequence or PDB file, or both. SEQ: sequence-based feature. STR: structure-based feature. DYN: dynamics-based feature.

| Feature Name | Description | Type |
| --- | --- | --- |
| Variation Number | Sequence position conservation score calculated using homologs<br>Variation numbers employed in the model are scaled using min-max normalization for each amino acid sequence | SEQ |
| $\Delta E$ Epistatic Score | Change in evolutionary statistical energy computed by EVmutation<sup>39</sup> | SEQ |
| Functional Impact Score (FIS) | Predicted magnitude of the effects of amino acid substitutions weighted by the relative frequency of disease-causing and neutral amino acid substitutions computed by FATHMM<sup>41</sup> | SEQ |
| $\Delta PSIC$ | Difference of PSIC scores for two amino acid residue variants computed by PolyPhen-2<sup>15</sup> | SEQ |
| Wild-type PSIC | PSIC score for wild type amino acid residue computed by PolyPhen-2<sup>15</sup> | SEQ |
| $\Delta \Delta G_{fold}$ | Folding free energy difference computed by FoldX<sup>43</sup> | STR |
| SASA | Solvent accessible surface area computed by FreeSASA<sup>44</sup> | STR |
| Mutant SSF | Knowledge-based potential for mutant amino acid variants computed by MAESTRO<sup>45</sup> | STR |
| Active Site Value | Calibrated probability of being a ligand-binding residue<br>Assigned 1 if the probability is greater than 0.5<br>computed by P2Rank<sup>46</sup> | STR |
| Mutant Reference Energy | Unfolded-state reference energies for mutant amino acid variants computed by PyRosetta<sup>47</sup> | STR |
| ΔReference Energy | Difference between unfolded-state reference energies for two amino acid variants computed by PyRosetta<sup>47</sup> | STR |
| MSD | Mean squared displacements of $C_\alpha$ atoms derived from the anisotropic network model computed by *ProDy*<sup>48</sup> | DYN |
| Mechanical Stiffness | Measurement of the mechanical resistance of residues to external pulling forces computed by *ProDy*<sup>48</sup> | DYN |
| Effectiveness | The ability of a residue to transmit mechanical deformation signals when subjected to a unit perturbation computed by *ProDy*<sup>49</sup> | DYN |
| Sensitivity | The ability of a residue to sense mechanical deformation signals when subjected to a unit perturbation computed by *ProDy*<sup>49</sup> | DYN |

**Figure 2.**
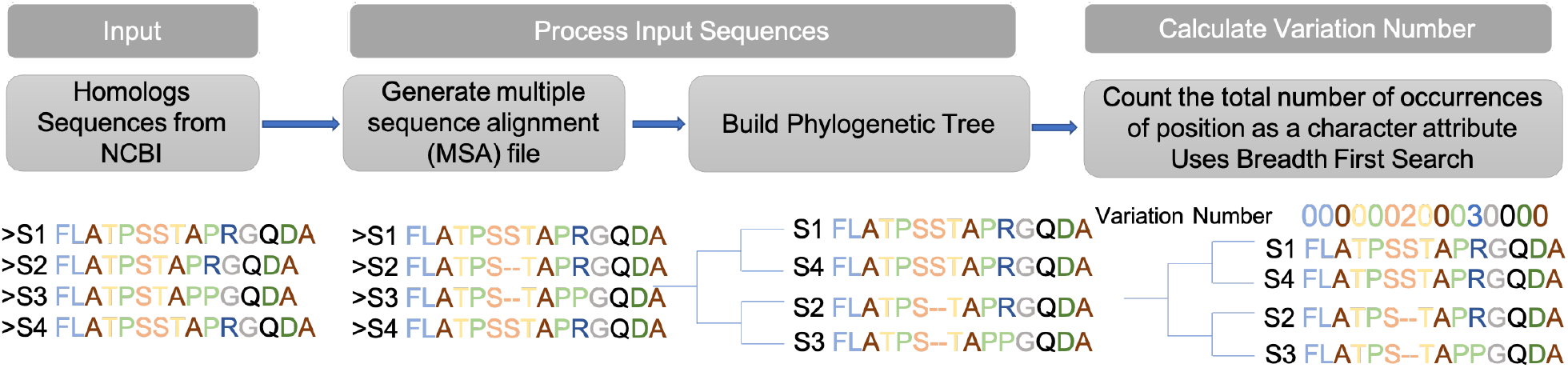
Pipeline for Calculating Variation Number. Variation number counts the number of occurrences of a position as a character attribute in all tree clades, where a character attribute may be defined as a state that exists in some elements of a clade, but not in the alternate clade under the same parent node^56^. For a given amino acid sequence, the homolog sequences are obtained using the NCBI Homologs Database. Multiple sequence alignment files are built using Clustal Omega^57^. Phylogenetic trees are generated using PAUP software with the maximum parsimony method^58^. Variation number, calculated using breadth first search, is the number of occurrences of a position as a character attribute in a given tree^36^. For each amino acid sequence, variation numbers at all of the human positions are normalized using min-max normalization. A smaller variation number suggests more conservation. The software is available at thesentence https://github.com/jiaying2508/variation-number.

Principal component analysis (PCA) is a way of identifying patterns in data that highlight their similarities and differences. The target dataset can be compressed by performing a PCA that reduces its number of dimensions if the data’s cumulative variance does not drop below a desired threshold, i.e., if there is not too much loss of information. Redundancy was analyzed for the 15 selected model features using PCA.

### Machine learning model selection and evaluation

TPOT was used in this study to determine the ML pipeline with the highest accuracy for our training dataset^52^. To prevent overfitting, we avoided any overlaps between training and evaluation datasets. Eighty percent of the dataset was used for training and twenty percent was used for testing. For TPOT parameters, both the number of generations and population size were set to 100, the cross validation size was set to 5, and the verbosity was set to 2.

TPOT suggested an XGBoost classifier to be the most suitable for our training dataset. The XGBoost algorithm, originally created by Chen and Guestrin, is a scalable tree boosting system that has been widely used by researchers^53^. Extensive studies have been done to showcase that XGBoost is very well-suited for building up a strong classification model, and it has been used to obtain many winning solutions in ML competitions^53–55^.

The performance of the chosen ML pipeline was compared to that of the PolyPhen-2, PROVEAN, SIFT, Rhapsody, EVMutation, MutationAssessor, SuSPect, FATHMM and MVP algorithms^15,34,37–42^. The performance of each method was assessed based upon its accuracy, sensitivity, specificity, F-measure, and Matthews Correlation Coefficient (MCC), as defined below:

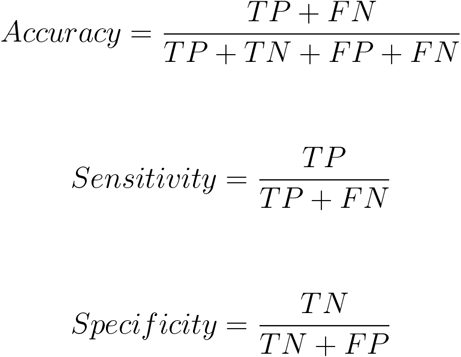

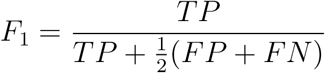

and

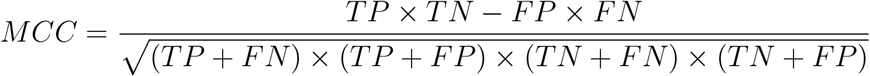

For those algorithms that could not predict the pathogenicity of an SAV, the corresponding SAV was left out of the performance assessments.

## Results

### Feature validation

Figure 3a shows the histograms of variation numbers for the pathogenic and neutral SAVs. Variation numbers range from 0 to 1, where 0 means high conservation and 1 means low conservation. The mean variation number for the pathogenic SAVs is 0.12, while the mean variation number for the neutral SAVs is 0.32. A t-test was performed using variation numbers for pathogenic and neutral SAVs^59^. The resulting t-statistic is −80.33, with a p-value of 0.0. The t-test results suggest that variation numbers for pathogenic and neutral SAVs are significantly different: pathogenic SAVs have smaller variation numbers than neutral SAVs, which suggests that pathogenic SAVs are more conserved than neutral SAVs.

**Figure 3.**
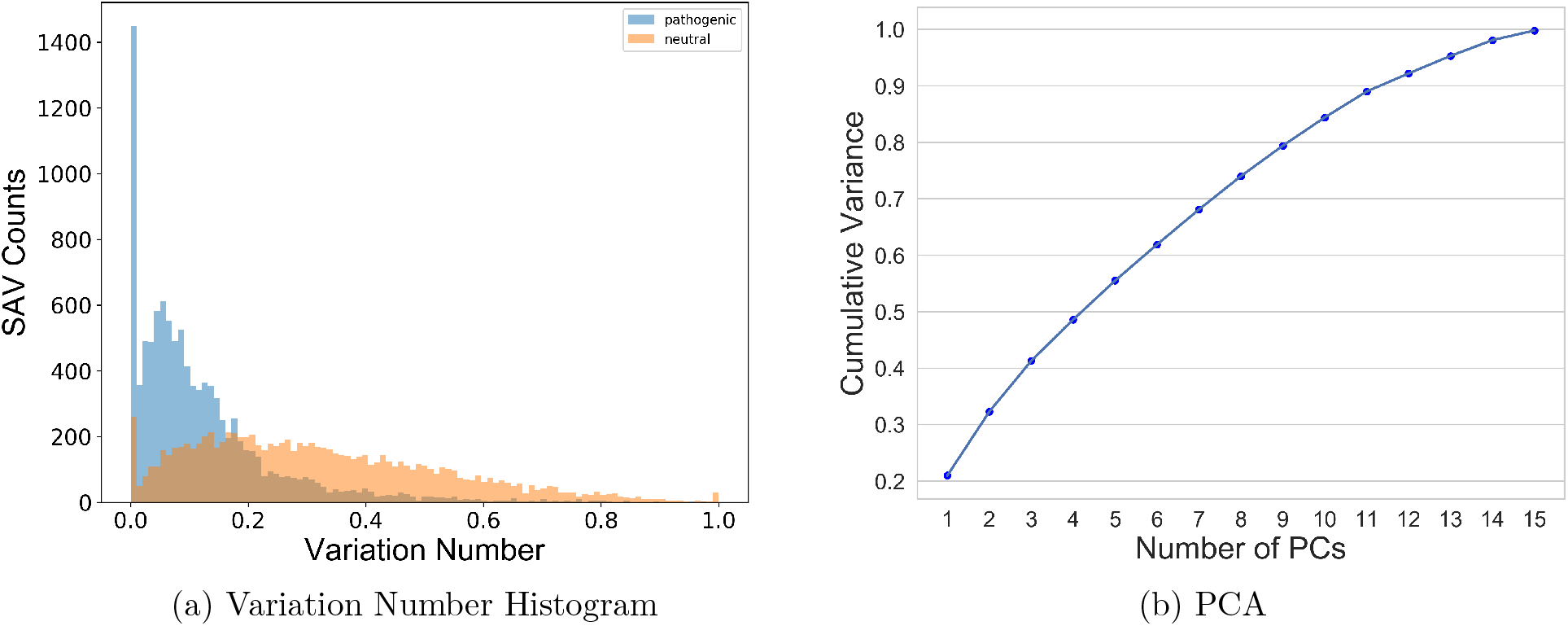
Feature Validations. (a) A comparison of variation number histograms for the pathogenic and neutral SAVs. Among the 20,307 selected SAVs, 9,743 SAVs were neutral and 10,564 SAVs were pathogenic. The mean variation number for the pathogenic SAVs was 0.12 while the mean variation number for the neutral SAVs was 0.32. (b) Plot of the cumulative variance vs. the number of principal components from a PCA analysis of our 15 features. This cumulative variance plot illustrates that 13 components are needed to describe 90% of the variance in the data.

PCA was applied to our feature dataset with the objective of cross-validating our feature selections and checking redundancy among our 15 features. Figure 3b shows the correlation between the cumulative variance (i.e., the sum of the variances of the individual principal components) and the number of principal components. The plot shows that 13 components are needed to describe 90% of the variance in the calculated results of all SAVs’ 15 features. Because most of the population variance cannot be attributed to the first few components, they cannot replace the original variables without loss of information. This analysis validates that there is minimal redundancy in our dataset and further supports the use of the selected features in the subsequently chosen ML model.

In addition to the PCA, Pearson correlations were calculated between all pairs of features, as depicted in Supplementary Figure S1. The top features that have the highest correlation with clinical scores are the wild-type PSIC, ΔPSIC, functional impact score, and variation number, which are all sequence-based features. Four pairs of features have a (negative) correlation greater than 0.5. The wild-type PSIC and ΔPSIC have a correlation of 0.66, the wild-type PSIC and variation number have a negative correlation of −0.59, the solvent accessible surface area (SASA) and mechanical stiffness have a negative correlation of - 0.53, and the mutant reference energy and mutant statistical scoring function (SSF) have a negative correlation of −0.52. All other pairs of features have (negative) correlations smaller than 0.5. The correlation heatmap of the raw data suggests that sequence features have a larger correlation with pathogenicity than the structural and dynamic features. It also shows that all of the features are largely independent of one another, and thus the inclusion of all of the features in our model is necessary.

### Machine learning pipeline

The classification model is intended for predicting whether an SAV is pathogenic (score 1) or non-pathogenic (score 0). TPOT recommended the XGBoost Classifier, which achieved the highest accuracy of 0.87, as the most suitable ML method for our dataset^52,53^. The optimized XGBoost classifier has a learning rate of 0.1. Feature importance scores were calculated and the average of 1000 repeats was taken(Supplementary Figure S2). ΔPSIC has the highest weight, followed by FIS, wild-type PSIC, and variation number, which are all sequence-based features. This is all in accordance with the feature correlation heatmap (Supplementary Figure S1). All of the other features had smaller, but similar importance values.

### Predictive power of the model

A total of 20,307 SAVs were extracted from ClinVar. To examine the performance of the model, 20% of the training dataset (4,062 SAVs) was randomly selected for testing purposes 1000 different times. Supplementary Table S2 present the true positive, true negative, false positive, and false negative counts for each method. Rhapsody and EVMutation are able to predict the correct pathogenicity of less than 60% of the SAVs. To further test the ability of the methods to distinguish pathogenic mutations, we plotted both the receiver operating characteristic (ROC) curve and the precision recall (PR) curve; the results are shown in Figures 4a and 4b. LYRUS has the highest AUC score of 0.941. It also dominates the PR space with a score of 0.95. The overall accuracy, sensitivity, specificity, F-measure (*F*_1_), and MCC were calculated for LYRUS as well as nine other methods (Supplementary Figure S3 and Table S3). LYRUS achieved the highest accuracy, specificity, F-measure, and MCC. LYRUS has a sensitivity of 0.890, which is lower than that of MVP (0.986), Polyphen-2 (0.924) and SIFT (0.906). These statistics demonstrate the power of our pipeline.

**Figure 4.**
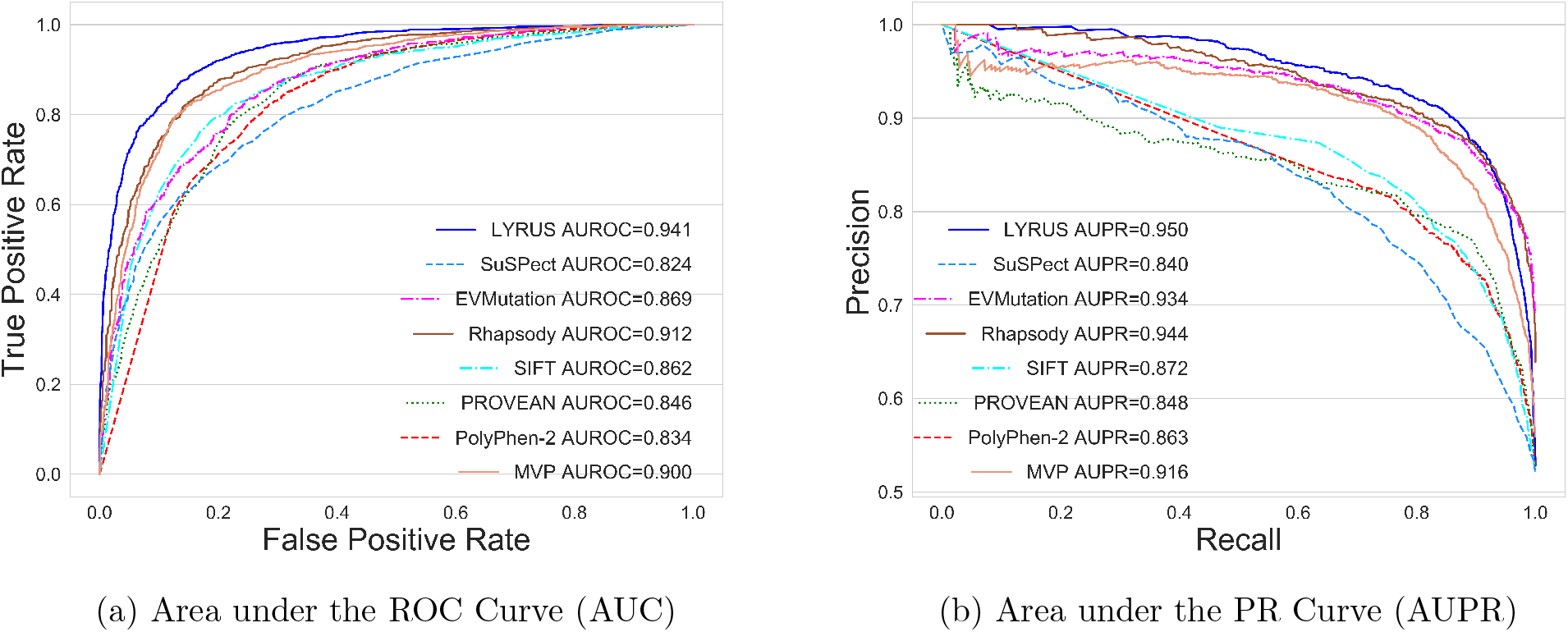
AUC and AUPR. (a) **Area under the ROC Curve (AUC).** The AUC of LYRUS was compared with that of SuSPect, EVMutation, Rhapsody, SIFT, PROVEAN, and PolyPhen2. The AUC for LYRUS was about 0.941, which was greater than that for all of the other methods studied here. (b) **Area under the PR Curve (AUPR).** The AUPR for LYRUS was 0.950, which was once again greater than that for all of the other methods.

## Illustrative Applications

To illustrate the effectiveness of LYRUS for identifying pathogenicity from neutral variants, we present a case study of two proteins, phosphatase and tensin homolog deleted on chromosome 10 (PTEN) and tumor protein 53 (TP53), which possess both pathogenic and neutral variants. Before being applied to PTEN and TP53, LYRUS was retrained on datasets that excluded the SAVs of these two proteins.

### Pathogenicity of PTEN mutants

PTEN is associated with advanced-stage or metastatic cancers^60–62^. LYRUS was applied to a dataset of 7,657 (403×19) SAVs of PTEN. PTEN (UNIPROT: P60484) has 403 amino acids. However, the complete X-ray crystal structure for PTEN is unavailable. The PDB 1D5R structure was used as a template to simulate PTEN using the Robetta server^63^. Simulated structures were used for the PTEN amino acids 1-13, 282-312, and 352-403, which are missing from the crystal structure. The prediction results for PTEN are shown in Supplementary Figure S4. Most SAVs in PTEN are predicted to be pathogenic, but all possible 342 mutants from Thr286 to Ser305 except Asn292 and Gly293 were predicted to be neutral. These positions are all located on the surface of the protein (Figure 5a), and the neutral predictions are due to their low ΔPSIC scores, negative or small positive ΔΔG_fold_ values, large SASA values, low stiffness values, and large MSD values.

**Figure 5.**
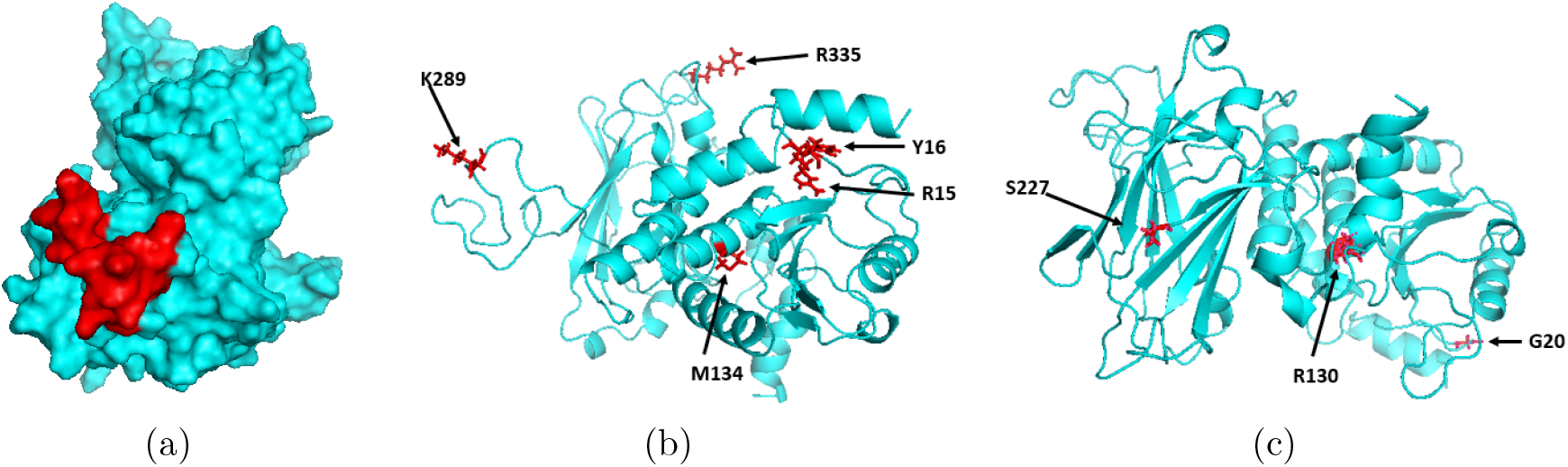
Cartoon and Surface Representations of the PTEN Protein. (a) The red surface corresponds to PTEN positions 286 to 305. (b) Visualization of the positions of the five false negative variants. (c) Visualization of the positions of the three false positive variants.

We used ClinVar and Humsavar to find PTEN SAVs with clinical implications^50,64^. There are 110 PTEN SAVs with a ‘review star’ of at least one in ClinVar^50^. Humsavar was also used to retrieve PTEN SAVs^64^. There are 52 SAVs that are categorized as either “Polymorphism” (equivalent to our score of 0) or “Disease” (equivalent to our score of 1). Between the two datasets, there are 29 repetitive SAVs, seven of which had contradicting annotations. All seven contradicting SAVs are classified as “Polymorphism” in Humsavar and “Pathogenic” in ClinVar. We used ClinVar annotations for these seven SAVs. In total, 133 SAVs for PTEN were used to test the performance of LYRUS against that of other methods. The results are listed in Supplementary Table S4 and Supplementary Figure S5. Only 39% of the SAVs can be predicted using EVMutation. SuSPect and FATHMM have slightly higher accuracies and sensitivities than LYRUS.

We further examine the SAVs whose pathogenicity is incorrectly predicted by LYRUS. There are five false negative SAVs: R15K, Y16H, R335Q, M134L, and K289E (Figure 5b). R15K is predicted to be neutral given its low ΔPSIC and WT PSIC values. ΔPSIC scores indicate the difference between the profile scores (obtained from computing the profile matrix^65^) of the two allelic variants in the polymorphic position^15^. Large positive values of this difference suggest that the studied substitution is rarely or never observed in the protein family. R15K’s small positive ΔPSIC value implies that this specific substitution is frequently observed in the protein family and hence less likely to be pathogenic^65^. The same rationale can be used to explain the remaining four false negative predictions (i.e., Y16H, R335Q, M134L, and K289E) even though their positive ΔΔG_fold_ values would suggest that they are destabilizing mutations. Interestingly, variation numbers of all five SAVs are relatively low, indicating that these five sites are highly conserved. This finding also demonstrates the efficacy of variation number in pathogenicity prediction. Despite these five false negative predictions, predictions based on dynamics-based features alone were largely correct. For example, it is evident from the PTEN structure (Figure 5b) that M134L belongs to a *β*-strand, which validates its small MSD and large stiffness values.

The three false positive SAVs are G20E, R130G, and S227F (Figure 5c). These three false positive SAVs are not solvent-exposed (Figure 5c), are buried within the structure, and accordingly form more interactions with neighboring residues. Interestingly, G20E is predicted to be pathogenic by all of the software studied (an EVMutation prediction is not available for G20E). Although G20E is annotated as a polymorphism in Humsavar, two studies suggested G20E to be a cancer-associated variant^66,67^, thus the annotation of G20E might be incorrect in Humsavar. R130G is labeled as a polymorphism in Humsavar. However, there were three other variants at position 130, R130Q, R130L, and R130P, that are recorded as pathogenic in ClinVar. R130G is predicted to be pathogenic, not only because it is located in the active site but also because of its calculated large ΔPSIC value and low variation number, ΔE, FIS, and SASA values. However, the computed ΔΔG_fold_ value of this variant is negative, which means that this SAV is presumably a stabilizing mutant or that it could be an overstabilizing mutant rendering it pathogenic. The last false positive is S227F whose incorrect prediction was due to its large ΔPSIC and ΔΔG_fold_ values, and low variation number, ΔE, and SASA values. Based on our calculations, the values of some features are closer to the averages of those of features for pathogenic SAVs and the rest are closer to the averages of the features for neutral SAVs. Therefore, inclusion of more relevant features (see Discussion) may prove to be an effective way to further improve the performance of LYRUS.

### Pathogenicity of TP53 mutants

LYRUS was also applied to TP53, which encodes a multifunction transcription factor whose loss promotes tumor formation^68^. The predicted probabilities of pathogenicity of the TP53 variants are presented in Supplementary Figure S6. The region spanning codons 100-290 is predicted to be highly pathogenic. This region contains the core domain of the TP53 protein, and mutations in the core domain can result in the loss of DNA binding activity^69^.

In addition, more than 80% of somatic TP53 mutations in human cancers occur in this region^69,70^. These findings validate our predictions. The performance of LYRUS was also compared with that of nine other software using 142 ClinVar entries (Supplementary Figure S7 and Supplementary Table S5). LYRUS achieved the second highest accuracy as well as sensitivity. SuSPect and FATHMM have much lower accuracy predicting the pathogenicity of SAVs in TP53 than in PTEN, highlighting the inconsistent performance of these two software. Furthermore, MVP achieved the highest sensitivity but the lowest specificity for both PTEN and TP53, substantiating the potential high false-negative rates. LYRUS, on the other hand, performed well in predicting both PTEN and TP53. There are four false positive predictions by LYRUS: Y107H, S185N, N235S, and G293W. All four SAVs are located on the surface of the protein and are hence solvent-exposed (Figure 6). They are all predicted to be pathogenic due to their high ΔPSIC and SASA values and low ΔE and FIS values.

**Figure 6.**
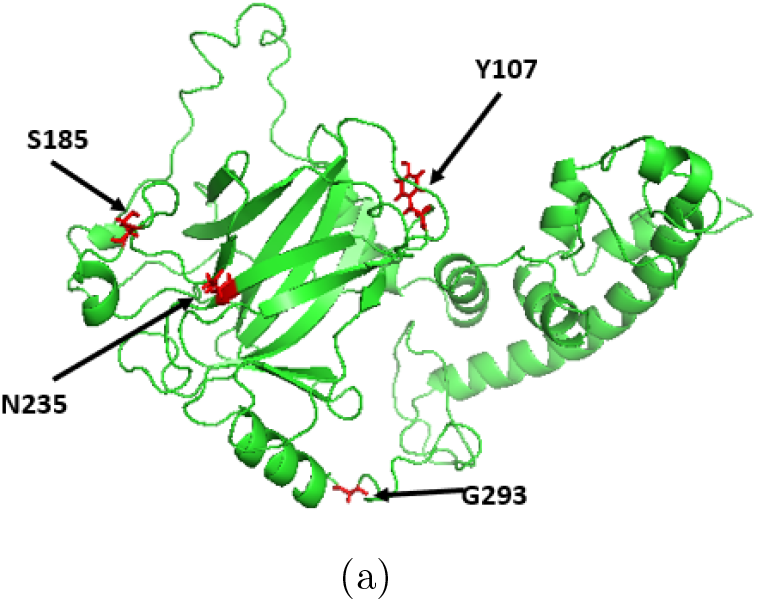
Carton Representation of the TP53 Protein. TP53 protein structure and the visualization of the positions of the false positive variants.

## Discussion

This study introduces LYRUS, an ML approach with the optimal pipeline selected by TPOT for predicting the pathogenicity of human SAVs. We aimed to develop an algorithm for predicting the clinical pathogenicity of human SAVs, thus the ClinVar database was used to generate the training dataset. Most methods in the field, such as PolyPhen2 and FATHMM, were designed to predict the effects of SAVs on protein function rather than their clinical significance^15,41^. Functional effects and clinical significance are not one and the same. However, in order to compare the predictions across a wider range of methods, we purposefully disregarded this subtlety.

Four pairs of features used by LYRUS have correlation coefficients higher than 0.5. The wild-type PSIC and ΔPSIC have a correlation of 0.66, which is expected since the model of sequence family evolution that computes the scores was constructed with the assumption that substitution probabilities are position-dependent^65^. The wild-type PSIC and variation number have a negative correlation of −0.59, which is intuitively reasonable considering that the lower the variation number, the more conserved a given amino acid is at a particular position, and the higher the PSIC score, the more likely this particular amino acid occurs at this position. SASA and mechanical stiffness have a negative correlation of −0.53, because buried residues with less solvent accessible surface area are more resistant to external pulling forces, thus exhibiting high mechanical stiffness.

The XGBoost classifier was picked by TPOT as the best model for our dataset. The XGBoost classifier minimizes data-overfitting issues^53^. A large number of false positives is often a consequence of overfitting, and by using the XGBoost classifier, this issue was minimized in LYRUS. LYRUS achieved the highest specificity among all the software to which we compared (Supplementary Figure S3 and Supplementary Table S3), while the sensitivity of LYRUS is lower than that of MVP, PolyPhen2, andSIFT. MVP achieved the highest sensitivity, suggesting the power of the method to identify all of the pathogenic variants. However, MVP has a very low specificity, which can be problematic as all the non-pathogenic variants in PTEN and TP53 are also classified as pathogenic (Supplementary Figure S5 and Supplementary Figure S7). LYRUS achieved the highest overall accuracy, F-measure, and MCC, demonstrating the software’s strong performance.

The most predictive features in LYRUS are sequence-based features. Studies have shown the importance of using amino acid conservation for pathogenicity prediction, which explains the high impact of sequence-based features in LYRUS^15,37,71^. The high impact score of variation number also suggests the effectiveness of this novel feature for categorizing pathogenic and non-pathogenic SAVs. Although structural and dynamics-based features have lower weights in LYRUS, these features are still valuable to include. Existing studies have shown that combining information gained from multiple sequence alignment and three-dimensional protein structures increases prediction performance^12,72^. Among the structural and dynamic features, change in folding free energies and the location of binding sites have the highest weights in LYRUS. In fact, several computational approaches have been developed to predict ΔΔG_fold_ in order to link it to the pathogenicity of mutations, which suggests the importance of incorporating ΔΔG_fold_ into LYRUS^19,20^. Catalytic residues, which comprise drug binding sites, are often conserved during evolution, and mutations of these residues can be detrimental^73^. This suggests the importance of incorporating information regarding the location of binding sites into pathogenicity predictors.

Although studies have demonstrated the utility of considering the equilibrium dynamics of proteins as a means of improving the predictive ability of pathogenicity predictors, our study reveals that dynamics-based features did not significantly contribute to the predictive power of LYRUS. One reason that dynamics-based features have a low impact score in LYRUS might be the limitations imposed by the models we used to calculate them. For example, the main disadvantage of the anisotropic network model (ANM) is its inability to account for anharmonic motions or multimeric transitions driven by a protein’s slowest collective modes^74^. The use of more sophisticated dynamics models may better capture the protein dynamics and further improve the prediction accuracy. The inclusion of other dynamic models are of future interest.

Another area for future improvement is the incorporation of structural changes caused by mutations into the model. LYRUS predicts the pathogenicity of SAVs based on the original protein structure instead of the mutated one. It has been proven that mutations promoting protein misfolding contribute to a variety of human diseases. Incorporating information related to structural changes, such as protein root mean square deviations (RMSDs), which reflect structural changes, may facilitate pathogenicity prediction^75–77^. Other thermodynamic information, such as changes in binding free energies, may also enhance the accuracy of the model. The prediction method may additionally be extended to other types of DNA mutations, such as insertions and deletions which may result in frameshifts.

SAVs from the Clinvar and Humsavar databases were used to test the performance of LYRUS on PTEN^50,64^. Twenty nine SAVs are present in both datasets, seven (24%) of which have contradictory annotations. The disagreement between the two datasets demonstrates the importance of choosing the most appropriate annotations for the SAVs.

In this study, a small part of the PTEN structure was simulated. However, because our method relies heavily on the PDB structure of the protein, we would not recommend applying LYRUS to a protein whose experimental PDB structure is unavailable. Thus, our method cannot be applied to proteins such as BRCA1, which is a limitation of our approach. With advances in protein folding algorithms, such as AlphaFold, it may become possible to predict the pathogenicity of SAVs using predicted structures^78^. Future work is needed to generate a pipeline which can be applied to simulated structures. LYRUS is built upon existing software (Supplementary Table S1), and the most computationally expensive part of the method is the calculation of ΔΔG_fold_ using FoldX. The current software is built to predict the SAVs for a single protein structure, and future improvement is needed to improve the computational efficiency and enable the prediction of SAVs from different protein structures.

## Conclusion

This study presents an ML pipeline (LYRUS) to predict human SAV pathogenicity that incorporates variation number along with 14 other features. LYRUS attained an accuracy of 0.87 using an autoML-selected (TPOT) XGBoost classifier. The XGBoost model suggests that sequence-based features have larger weights than structural and dynamic features in SAV pathogenicity prediction. Variation number feature is negatively correlated with clinical pathogenicity, and has the fourth highest weight among all of the features studied here. Comparisons among LYRUS and nine other methods suggested that our model had the highest overall accuracy, specificity, F-measure, and MCC. Application to PTEN and TP53 also showed the high and consistent performance of LYRUS compared to other software. The scripts for LYRUS are available at https://github.com/jiaying2508/LYRUS.

## Acknowledgements

J.L. and I.S. thank Daniel Weinreich and Sorin Istrail for discussions regarding the broad ideas underpinning this work. J.Y. and B.R. thank Marty Ytreberg, Jagdish Patel, and Daniel Weinreich for inspiring conversations and financial support. The authors would like to thank the Brown Center for Biomedical Informatics (BCBI) and the Brown Center for Computation and Visualization (CCV) for computing resources.

## Supplemental Information

**Figure S1.**
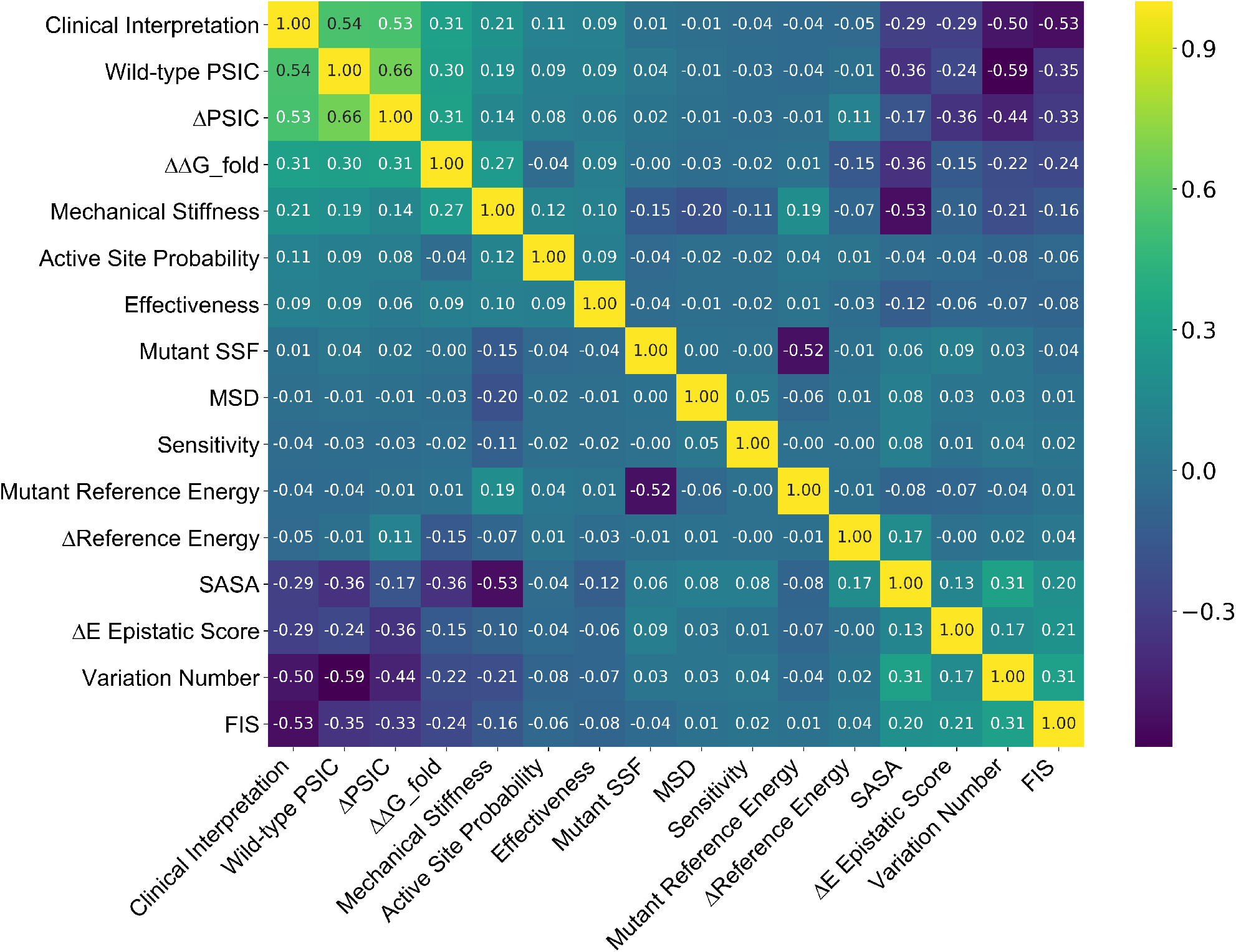
Feature Correlation Heatmap. Heatmap of the Pearson correlation coefficient for each pair of the features. The features with the highest correlation with the ClinVar scores were all sequence-based features. Of the 105 possible pairs of features, only four pairs had a (negative) correlation coefficient greater than 0.5, suggesting that the selected features were largely independent.

**Figure S2.**
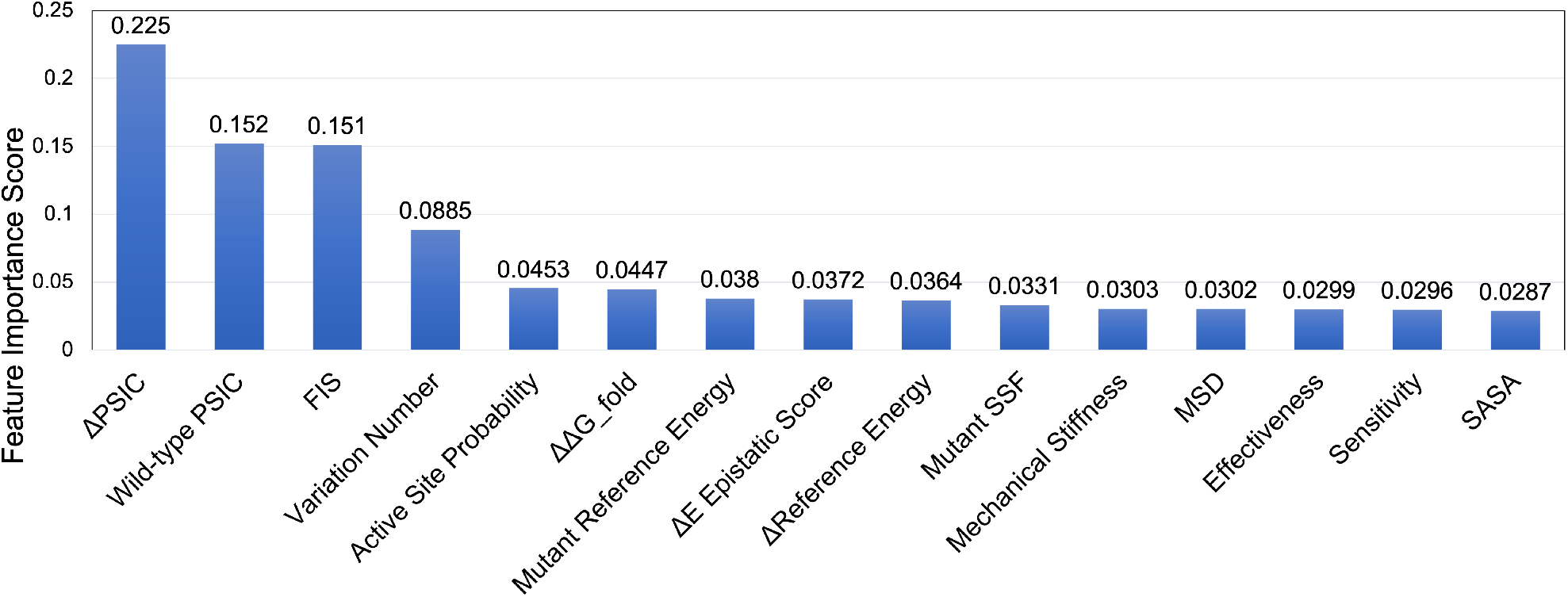
Feature Importance Scores from LYRUS. The sequence-based features had higher weights than structural and dynamics-based features. ΔPSIC had the highest importance score, followed by the wild-type PSIC, FIS, and variation number. The remaining 11 features had similar importance scores.

**Figure S3.**
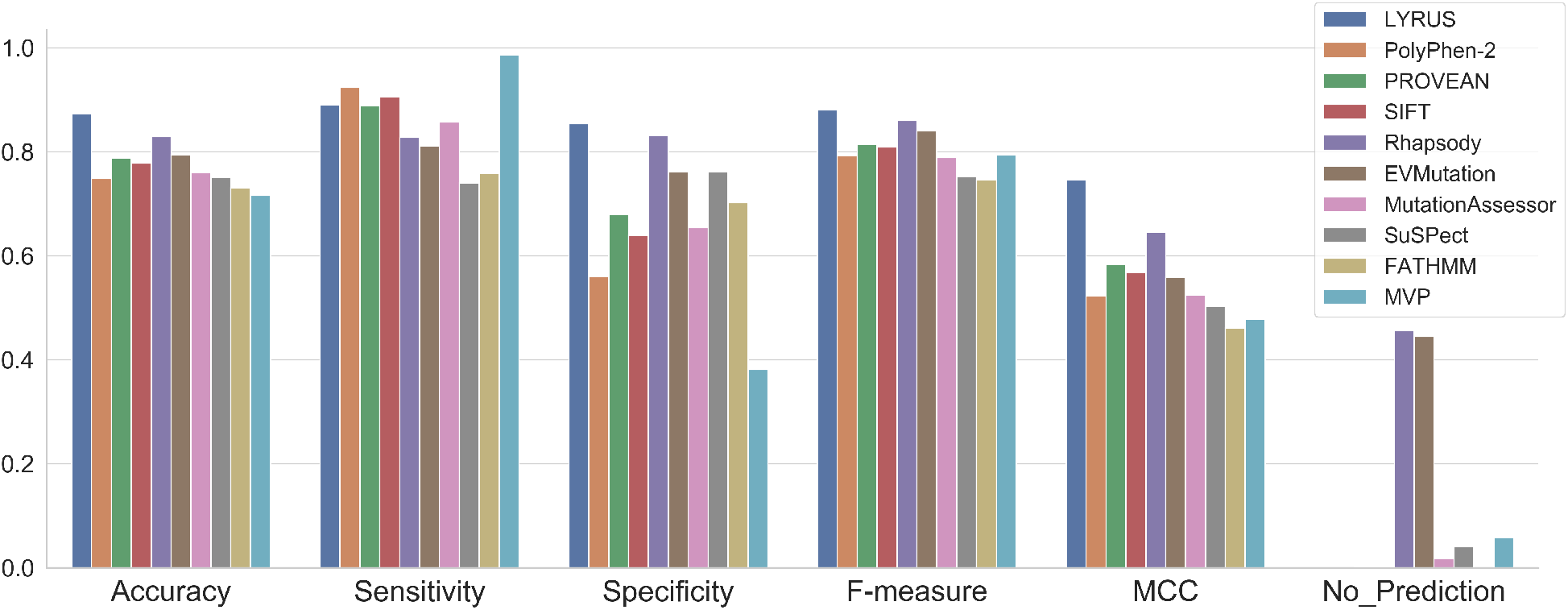
Statistics Compared to Other Software. The accuracy, sensitivity, specificity, F-measure, and MCC for each of the prediction methods were calculated. LYRUS achieved the greatest accuracy, specificity, F-measure, and MCC. The sensitivity of LYRUS was lower than that of MVP, PolyPhen2, and SIFT. Rhapsody and EVMutation were able to predict less than 60% of the the SAVs.

**Figure S4.**
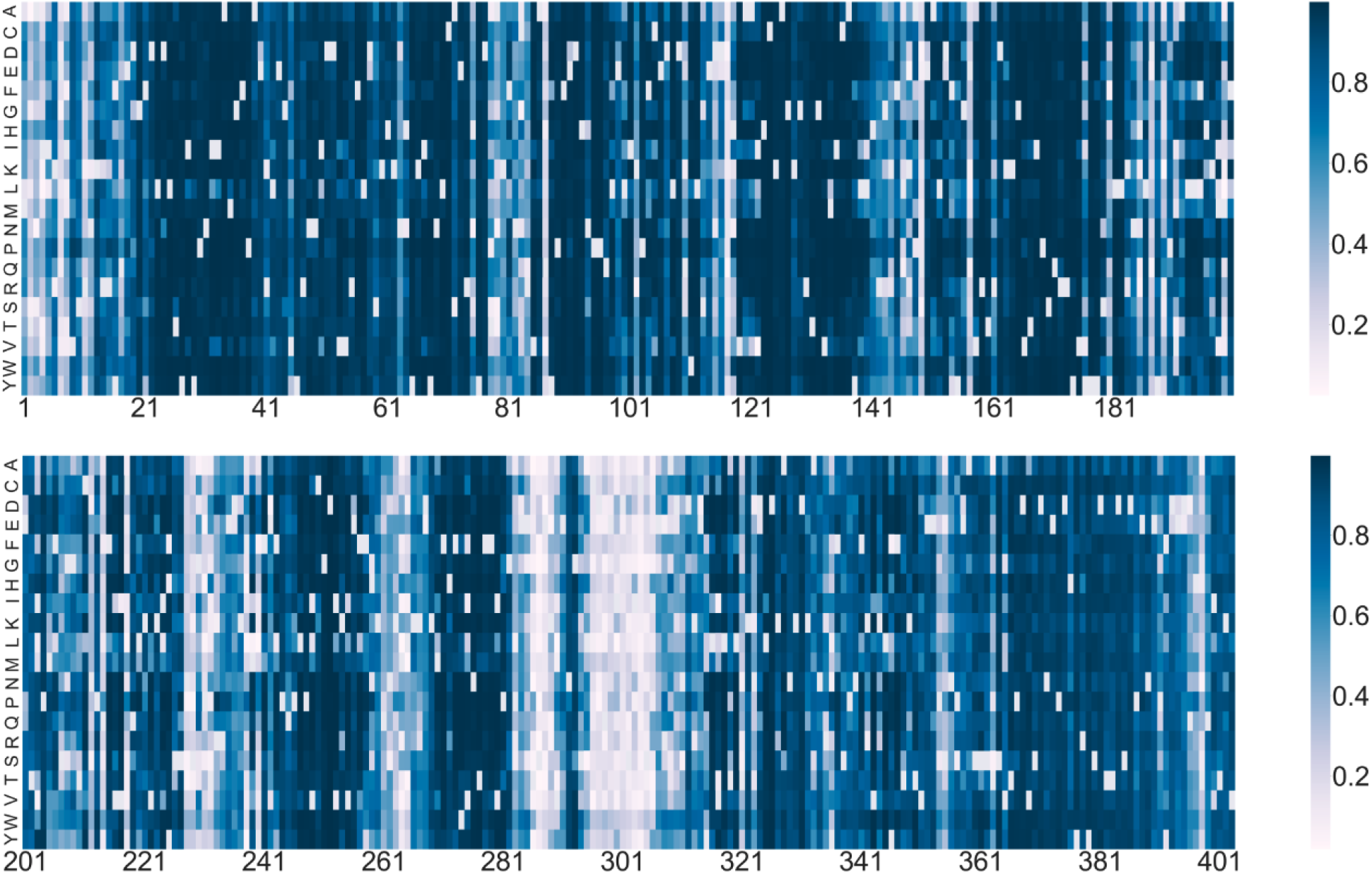
PTEN Prediction Heatmap. The x-axis represents PTEN amino acid positions and the y-axis represents different amino acid substitutions. The color coding of each heatmap cell represents the predicted probability of the SAV being pathogenic. Wild-type amino acids were assigned a probability of 0. LYRUS predicts most PTEN SAVs to be pathogenic.

**Figure S5.**
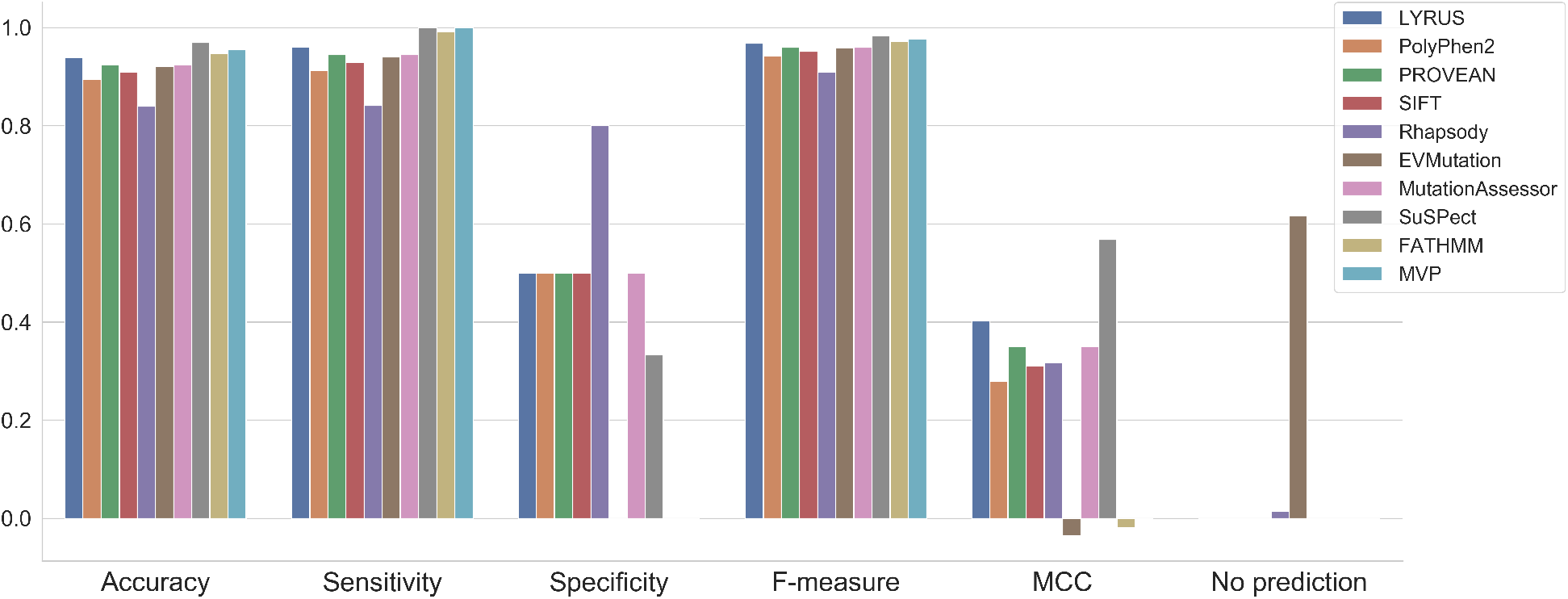
PTEN Statistics Compared to Other Software. The accuracy, sensitivity, specificity, F-measure, and MCC of LYRUS, PolyPhen2, PROVEAN, SIFT, Rhapsody, EVMutation, MutationAssessor, SuSPect, FATHMM, and MVP.

**Figure S6.**
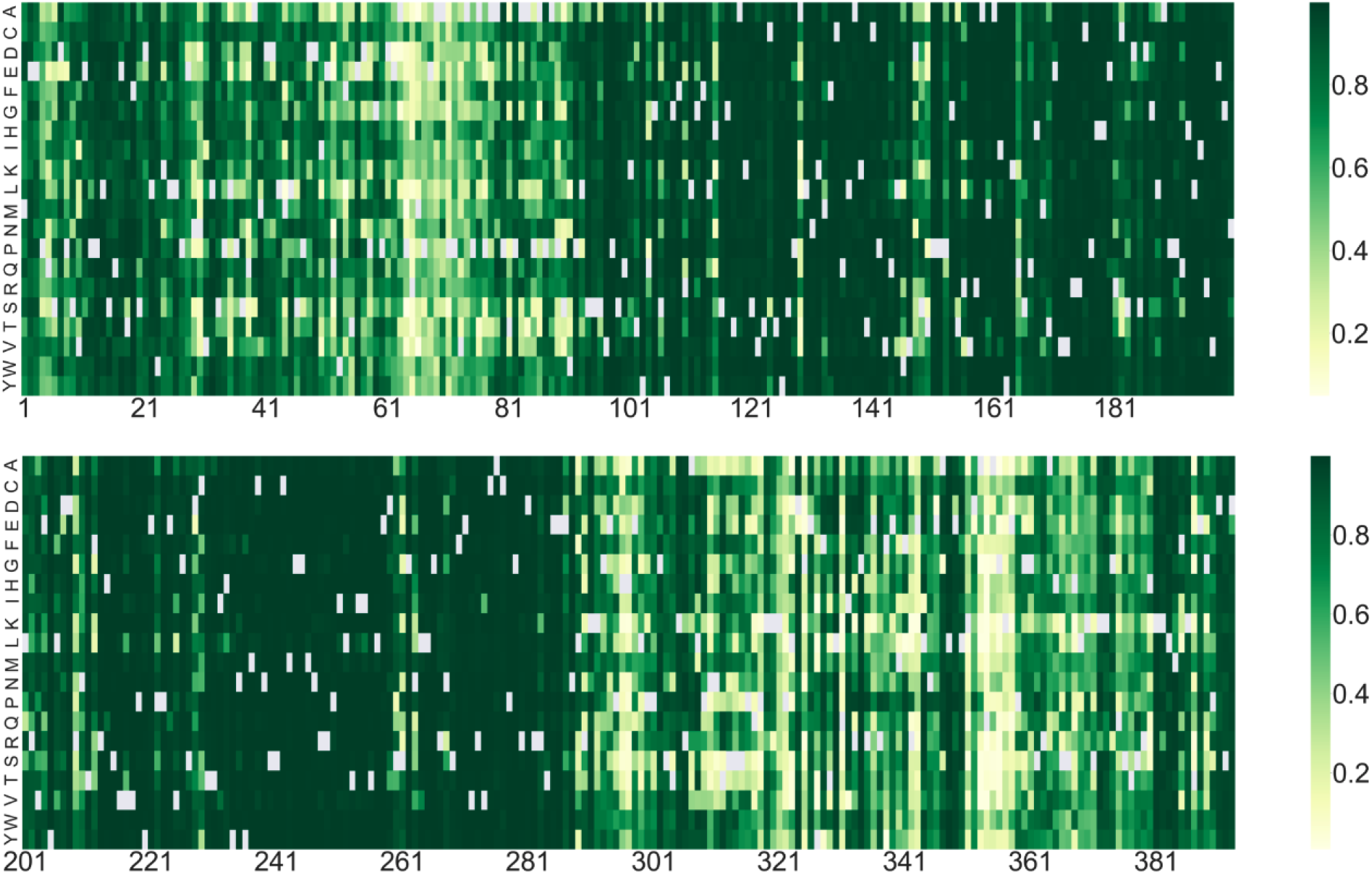
TP53 Prediction Heatmap. The x-axis represents TP53 amino acid positions and the y-axis represents different amino acid substitutions. The color coding of each heatmap cell represents the predicted probability of the SAV being pathogenic. Wild-type amino acids were assigned a probability of 0.

**Figure S7.**
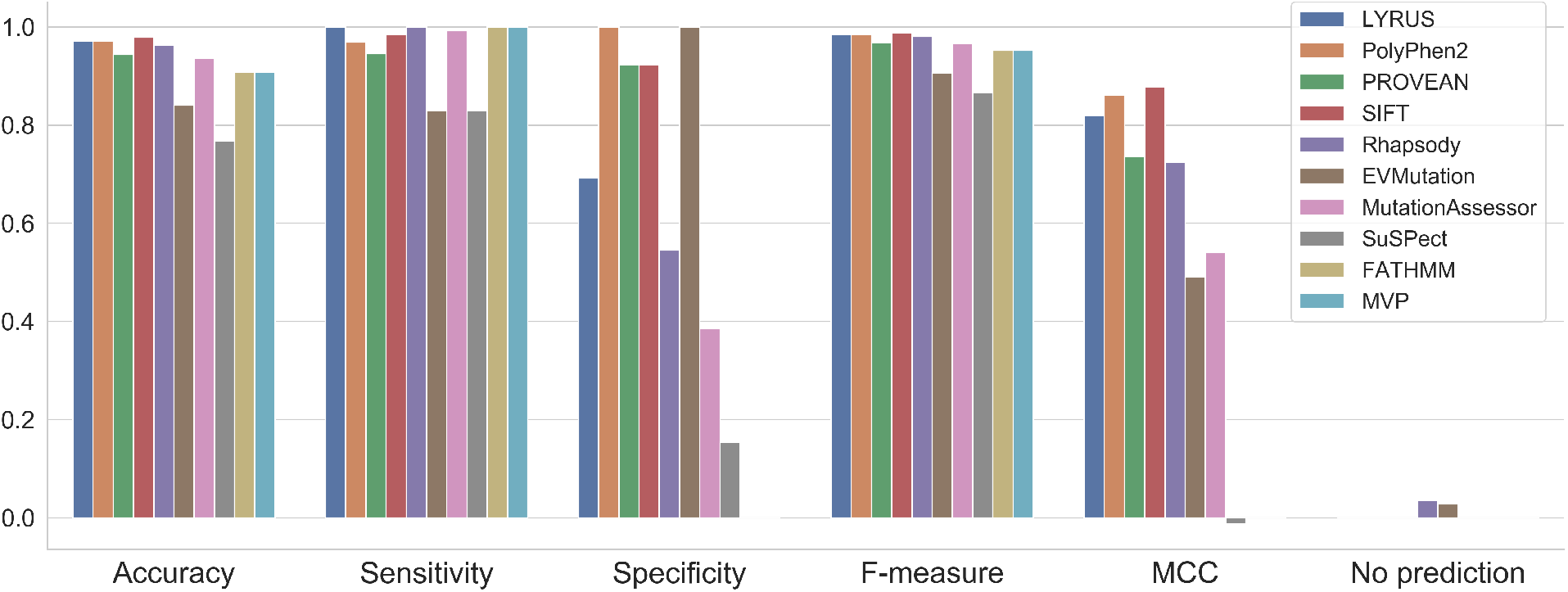
TP53 Statistics Compared to Other Software. The accuracy, sensitivity, specificity, F-measure and MCC of LYRUS, PolyPhen2, PROVEAN, SIFT, Rhapsody, EVMutation, MutationAssessor, SuSPect, FATHMM, and MVP.

**Table S1.** Links to the software used to compute each feature.

| Feature Name | Link |
| --- | --- |
| Variation Number | https://github.com/jiaying2508/XGBClassifier |
| $\Delta E$ Epistatic Score | https://github.com/debbiemarkslab/EVmutation |
| FIS | http://fathmm.biocompute.org.uk |
| $\Delta PSIC$ | http://genetics.bwh.harvard.edu/pph2/ |
| Wild-type PSIC | http://genetics.bwh.harvard.edu/pph2/ |
| $\Delta\Delta G_{fold}$ | http://foldxsuite.crg.eu |
| SASA | https://freesasa.github.io |
| Mutant SSF | https://pbwww.che.sbg.ac.at/?page_id=416 |
| Active Site Value | https://github.com/rdk/p2rank |
| Mutant Reference Energy | http://www.pyrosetta.org |
| ΔReference Energy | http://www.pyrosetta.org |
| MSD | http://prody.csb.pitt.edu |
| Mechanical Stiffness | http://prody.csb.pitt.edu |
| Effectiveness | http://prody.csb.pitt.edu |
| Sensitivity | http://prody.csb.pitt.edu |

**Table S2.**
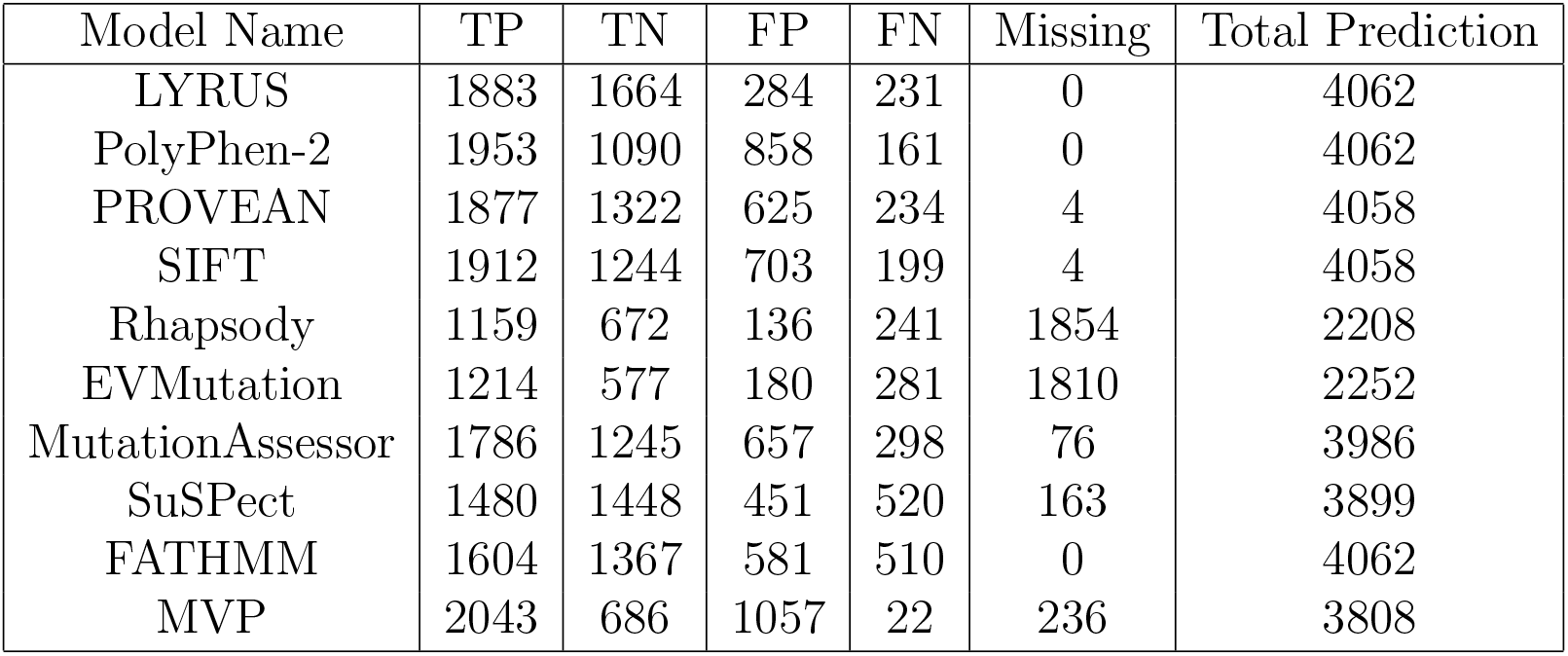
Comparison to Other Models. TP = True Positive. TN = True Negative. FP = False Positive. FN = False Negative.

**Table S3.** Comparison to Other Models. MCC =Matthews Correlation Coefficient.

| Model Name | Accuracy | Sensitivity | Specificity | F-measure | MCC | No Prediction |
| --- | --- | --- | --- | --- | --- | --- |
| LYRUS | 0.873 | 0.890 | 0.854 | 0.880 | 0.746 | 0% |
| PolyPhen-2 | 0.749 | 0.924 | 0.560 | 0.793 | 0.523 | 0% |
| PROVEAN | 0.788 | 0.889 | 0.679 | 0.814 | 0.583 | 0.101% |
| SIFT | 0.778 | 0.906 | 0.639 | 0.809 | 0.568 | 0.101% |
| Rhapsody | 0.829 | 0.828 | 0.832 | 0.860 | 0.645 | 45.644% |
| EVMutation | 0.795 | 0.812 | 0.762 | 0.840 | 0.559 | 44.565% |
| MutationAssessor | 0.760 | 0.857 | 0.654 | 0.789 | 0.525 | 1.864% |
| SuSPect | 0.751 | 0.740 | 0.762 | 0.753 | 0.503 | 4.003% |
| FATHMM | 0.731 | 0.759 | 0.702 | 0.746 | 0.461 | 0% |
| MVP | 0.716 | 0.986 | 0.381 | 0.794 | 0.478 | 5.810% |

**Table S4.** PTEN Case Study. A total of 133 SAV classifications are available from ClinVar and Humsavar. True positive (TP), true negative (TN), false positive (FP). false negative (FN), no prediction (NP), accuracy, sensitivity, specificity, F-measure and MCC are listed.

| Model Name | TP | TN | FP | FN | NP | Accuracy | Sensitivity | Specificity | F-measure | MCC |
| --- | --- | --- | --- | --- | --- | --- | --- | --- | --- | --- |
| LYRUS | 122 | 3 | 3 | 5 | 0 | 0.940 | 0.961 | 0.5 | 0.968 | 0.402 |
| PolyPhen-2 | 116 | 3 | 3 | 11 | 0 | 0.895 | 0.913 | 0.5 | 0.943 | 0.280 |
| PROVEAN | 120 | 3 | 3 | 7 | 0 | 0.925 | 0.945 | 0.5 | 0.96 | 0.350 |
| SIFT | 118 | 3 | 3 | 9 | 0 | 0.910 | 0.929 | 0.5 | 0.952 | 0.311 |
| Rhapsody | 106 | 4 | 1 | 20 | 2 | 0.840 | 0.841 | 0.8 | 0.910 | 0.318 |
| EVMutation | 47 | 0 | 1 | 3 | 82 | 0.922 | 0.94 | 0 | 0.959 | -0.035 |
| MutationAssessor | 120 | 3 | 3 | 7 | 0 | 0.925 | 0.945 | 0.5 | 0.96 | 0.350 |
| SuSPect | 127 | 2 | 4 | 0 | 0 | 0.970 | 1 | 0.333 | 0.984 | 0.568 |
| FATHMM | 126 | 0 | 6 | 1 | 0 | 0.947 | 0.992 | 0 | 0.973 | -0.019 |
| MVP | 127 | 0 | 6 | 0 | 0 | 0.955 | 1 | 0 | 0.977 | N/A |

**Table S5.** TP53 Case Study. A total of 142 SAV classifications are available from ClinVar. True positive (TP), true negative (TN), false positive (FP). false negative (FN), no prediction (NP), accuracy, sensitivity, specificity, F-measure and MCC are listed.

| Model Name | TP | TN | FP | FN | NP | Accuracy | Sensitivity | Specificity | F-measure | MCC |
| --- | --- | --- | --- | --- | --- | --- | --- | --- | --- | --- |
| LYRUS | 129 | 9 | 4 | 0 | 0 | 0.972 | 1.0 | 0.692 | 0.985 | 0.819 |
| PolyPhen-2 | 125 | 13 | 0 | 4 | 0 | 0.972 | 0.969 | 1.0 | 0.984 | 0.861 |
| PROVEAN | 122 | 12 | 1 | 7 | 0 | 0.944 | 0.946 | 0.923 | 0.968 | 0.736 |
| SIFT | 127 | 12 | 1 | 2 | 0 | 0.979 | 0.984 | 0.923 | 0.988 | 0.878 |
| Rhapsody | 126 | 6 | 5 | 0 | 5 | 0.964 | 1.0 | 0.545 | 0.981 | 0.724 |
| EVMutation | 107 | 9 | 0 | 22 | 4 | 0.841 | 0.829 | 1.0 | 0.907 | 0.491 |
| MutationAssessor | 128 | 5 | 8 | 1 | 0 | 0.937 | 0.992 | 0.385 | 0.966 | 0.540 |
| SuSPect | 107 | 2 | 11 | 22 | 0 | 0.768 | 0.829 | 0.154 | 0.866 | -0.013 |
| FATHMM | 129 | 0 | 13 | 0 | 0 | 0.908 | 1.0 | 0 | 0.952 | N/A |
| MVP | 129 | 0 | 13 | 0 | 0 | 0.908 | 1.0 | 0 | 0.952 | N/A |

## References

(1) Fowler, D. M.; Fields, S. Deep mutational scanning: a new style of protein science. Nature methods 2014, 11, 801–807.

(2) Shendure, J.; Balasubramanian, S.; Church, G. M.; Gilbert, W.; Rogers, J.; Schloss, J. A.; Waterston, R. H. DNA sequencing at 40: past, present and future. Nature 2017, 550, 345–353.

(3) Ormond, K. E.; Wheeler, M. T.; Hudgins, L.; Klein, T. E.; Butte, A. J.; Altman, R. B.; Ashley, E. A.; Greely, H. T. Challenges in the clinical application of whole-genome sequencing. The Lancet 2010, 375, 1749–1751.

(4) Ramensky, V.; Bork, P.; Sunyaev, S. Human non-synonymous SNPs: server and survey. Nucleic Acids Research 2002, 30, 3894–3900.

(5) Syvänen, A.-C. Accessing genetic variation: genotyping single nucleotide polymorphisms. Nature Reviews Genetics 2001, 2, 930–942.

(6) Cargill, M.; Altshuler, D.; Ireland, J.; Sklar, P.; Ardlie, K.; Patil, N.; Lane, C. R.; Lim, E. P.; Kalyanaraman, N.; Nemesh, J., et al. Characterization of single-nucleotide polymorphisms in coding regions of human genes. Nature genetics 1999, 22, 231–238.

(7) Yip, Y. L.; Scheib, H.; Diemand, A. V.; Gattiker, A.; Famiglietti, L. M.; Gasteiger, E.; Bairoch, A. The Swiss-Prot variant page and the ModSNP database: a resource for sequence and structure information on human protein variants. Human mutation 2004, 23, 464–470.

(8) Marinko, J. T.; Huang, H.; Penn, W. D.; Capra, J. A.; Schlebach, J. P.; Sanders, C. R. Folding and misfolding of human membrane proteins in health and disease: from single molecules to cellular proteostasis. Chemical reviews 2019, 119, 5537–5606.

(9) Yip, Y. L.; Famiglietti, M.; Gos, A.; Duek, P. D.; David, F. P.; Gateau, A.; Bairoch, A. Annotating single amino acid polymorphisms in the UniProt/Swiss-Prot knowledgebase. Human mutation 2008, 29, 361–366.

(10) Niu, B.; Scott, A. D.; Sengupta, S.; Bailey, M. H.; Batra, P.; Ning, J.; Wyczalkowski, M. A.; Liang, W.-W.; Zhang, Q.; McLellan, M. D., et al. Protein-structure-guided discovery of functional mutations across 19 cancer types. Nature genetics 2016, 48, 827–837.

(11) Yue, P.; Li, Z.; Moult, J. Loss of protein structure stability as a major causative factor in monogenic disease. Journal of molecular biology 2005, 353, 459–473.

(12) Saunders, C. T.; Baker, D. Evaluation of structural and evolutionary contributions to deleterious mutation prediction. Journal of molecular biology 2002, 322, 891–901.

(13) Sunyaev, S.; Ramensky, V.; Bork, P. Towards a structural basis of human non-synonymous single nucleotide polymorphisms. Trends in Genetics 2000, 16, 198–200.

(14) Wang, Z.; Moult, J. SNPs, protein structure, and disease. Human mutation 2001, 17, 263–270.

(15) Adzhubei, I. A.; Schmidt, S.; Peshkin, L.; Ramensky, V. E.; Gerasimova, A.; Bork, P.; Kondrashov, A. S.; Sunyaev, S. R. A method and server for predicting damaging missense mutations. Nature methods 2010, 7, 248–249.

(16) Capriotti, E.; Altman, R. B. Improving the prediction of disease-related variants using protein three-dimensional structure. BMC bioinformatics 2011, 12, S3.

(17) Ancien, F.; Pucci, F.; Godfroid, M.; Rooman, M. Prediction and interpretation of deleterious coding variants in terms of protein structural stability. Scientific reports 2018, 8, 1–11.

(18) Joerger, A. C.; Fersht, A. R. Structural biology of the tumor suppressor p53 and cancer-associated mutants. Advances in cancer research 2007, 97, 1–23.

(19) Peng, Y.; Alexov, E. Investigating the linkage between disease-causing amino acid variants and their effect on protein stability and binding. Proteins: Structure, Function, and Bioinformatics 2016, 84, 232–239.

(20) Petukh, M.; Kucukkal, T. G.; Alexov, E. On human disease-causing amino acid variants: Statistical study of sequence and structural patterns. Human mutation 2015, 36, 524–534.

(21) Blanco, J. D.; Radusky, L.; Climente-González, H.; Serrano, L. FoldX accurate structural protein–DNA binding prediction using PADA1 (Protein Assisted DNA Assembly 1). Nucleic acids research 2018, 46, 3852–3863.

(22) Zhang, Z.; Wang, L.; Gao, Y.; Zhang, J.; Zhenirovskyy, M.; Alexov, E. Predicting folding free energy changes upon single point mutations. Bioinformatics 2012, 28, 664–671.

(23) Getov, I.; Petukh, M.; Alexov, E. SAAFEC: predicting the effect of single point mutations on protein folding free energy using a knowledge-modified MM/PBSA approach. International journal of molecular sciences 2016, 17, 512.

(24) Li, M.; Petukh, M.; Alexov, E.; Panchenko, A. R. Predicting the impact of missense mutations on protein–protein binding affinity. Journal of chemical theory and computation 2014, 10, 1770–1780.

(25) Cang, Z.; Wei, G.-W. Analysis and prediction of protein folding energy changes upon mutation by element specific persistent homology. Bioinformatics 2017, 33, 3549–3557.

(26) Cang, Z.; Wei, G.-W. TopologyNet: Topology based deep convolutional and multi-task neural networks for biomolecular property predictions. PLoS computational biology 2017, 13, e1005690.

(27) Cheng, T. M.; Lu, Y.-E.; Vendruscolo, M.; Blundell, T. L., et al. Prediction by graph theoretic measures of structural effects in proteins arising from non-synonymous single nucleotide polymorphisms. PLoS Comput Biol 2008, 4, e1000135.

(28) Bao, L.; Cui, Y. Prediction of the phenotypic effects of non-synonymous single nucleotide polymorphisms using structural and evolutionary information. Bioinformatics 2005, 21, 2185–2190.

(29) Vendruscolo, M.; Dokholyan, N. V.; Paci, E.; Karplus, M. Small-world view of the amino acids that play a key role in protein folding. Physical Review E 2002, 65, 061910.

(30) Jacobs, D. J.; Rader, A. J.; Kuhn, L. A.; Thorpe, M. F. Protein flexibility predictions using graph theory. Proteins: Structure, Function, and Bioinformatics 2001, 44, 150–165.

(31) Kannan, N.; Vishveshwara, S. Identification of side-chain clusters in protein structures by a graph spectral method. Journal of molecular biology 1999, 292, 441–464.

(32) Ponzoni, L.; Bahar, I. Structural dynamics is a determinant of the functional significance of missense variants. Proceedings of the National Academy of Sciences 2018, 115, 4164–4169.

(33) Smith, I. N.; Thacker, S.; Seyfi, M.; Cheng, F.; Eng, C. Conformational dynamics and allosteric regulation landscapes of germline PTEN mutations associated with autism compared to those associated with cancer. The American Journal of Human Genetics 2019, 104, 861–878.

(34) Ponzoni, L.; Peñaherrera, D. A.; Oltvai, Z. N.; Bahar, I. Rhapsody: predicting the pathogenicity of human missense variants. Bioinformatics 2020, 36, 3084–3092.

(35) Banzhaf, W.; Nordin, P.; Keller, R.; Francone, F. GP–An Introduction; On the Automatic Evolution of Computer Programs and its Applications. 1998.

(36) Lai, J.; Sarkar, I. N. A Phylogenetic Approach to Analyze the Conservativeness of BRCA1 and BRCA2 Mutations. AMIA Annual Symposium Proceedings 2020,

(37) Choi, Y.; Sims, G. E.; Murphy, S.; Miller, J. R.; Chan, A. P. Predicting the functional effect of amino acid substitutions and indels. PLoS One 2012, 7, e46688.

(38) Kumar, P.; Henikoff, S.; Ng, P. C. Predicting the effects of coding non-synonymous variants on protein function using the SIFT algorithm. Nat. Protoc. 2009, 4, 1073–1081.

(39) Hopf, T. A.; Ingraham, J. B.; Poelwijk, F. J.; Schärfe, C. P. I.; Springer, M.; Sander, C.; Marks, D. S. Mutation effects predicted from sequence co-variation. Nat. Biotechnol. 2017, 35, 128–135.

(40) Reva, B.; Antipin, Y.; Sander, C. Predicting the functional impact of protein mutations: application to cancer genomics. Nucleic Acids Res. 2011, 39, e118.

(41) Shihab, H. A.; Gough, J.; Cooper, D. N.; Stenson, P. D.; Barker, G. L. A.; Edwards, K. J.; Day, I. N. M.; Gaunt, T. R. Predicting the functional, molecular, and phenotypic consequences of amino acid substitutions using hidden Markov models. Hum. Mutat. 2013, 34, 57–65.

(42) Qi, H.; Zhang, H.; Zhao, Y.; Chen, C.; Long, J. J.; Chung, W. K.; Guan, Y.; Shen, Y. MVP predicts the pathogenicity of missense variants by deep learning. Nat. Commun. 2021, 12, 510.

(43) Schymkowitz, J.; Borg, J.; Stricher, F.; Nys, R.; Rousseau, F.; Serrano, L. The FoldX web server: an online force field. Nucleic acids research 2005, 33, W382–W388.

(44) Mitternacht, S. FreeSASA: An open source C library for solvent accessible surface area calculations. F1000Research 2016, 5.

(45) Laimer, J.; Hofer, H.; Fritz, M.; Wegenkittl, S.; Lackner, P. MAESTRO-multi agent stability prediction upon point mutations. BMC bioinformatics 2015, 16, 116.

(46) Krivák, R.; Hoksza, D. P2Rank: machine learning based tool for rapid and accurate prediction of ligand binding sites from protein structure. Journal of cheminformatics 2018, 10, 39.

(47) Alford, R. F.; Leaver-Fay, A.; Jeliazkov, J. R.; O’Meara, M. J.; DiMaio, F. P.; Park, H.; Shapovalov, M. V.; Renfrew, P. D.; Mulligan, V. K.; Kappel, K., et al. The Rosetta all-atom energy function for macromolecular modeling and design. Journal of chemical theory and computation 2017, 13, 3031–3048.

(48) Bakan, A.; Meireles, L. M.; Bahar, I. ProDy: protein dynamics inferred from theory and experiments. Bioinformatics 2011, 27, 1575–1577.

(49) General, I. J.; Liu, Y.; Blackburn, M. E.; Mao, W.; Gierasch, L. M.; Bahar, I. ATPase subdomain IA is a mediator of interdomain allostery in Hsp70 molecular chaperones. PLoS Comput Biol 2014, 10, e1003624.

(50) Landrum, M. J. et al. ClinVar: improving access to variant interpretations and supporting evidence. Nucleic Acids Res. 2018, 46, D1062–D1067.

(51) Kiefer, F.; Arnold, K.; Künzli, M.; Bordoli, L.; Schwede, T. The SWISS-MODEL Repository and associated resources. Nucleic Acids Research 2008, 37, D387–D392.

(52) Le, T. T.; Fu, W.; Moore, J. H. Scaling tree-based automated machine learning to biomedical big data with a feature set selector. Bioinformatics 2020, 36, 250–256.

(53) Chen, T.; Guestrin, C. XGBoost: A Scalable Tree Boosting System. Proceedings of the 22nd ACM SIGKDD International Conference on Knowledge Discovery and Data Mining. New York, NY, USA, 2016; pp 785–794.

(54) Dhaliwal, S. S.; Nahid, A.-A.; Abbas, R. Effective intrusion detection system using XGBoost. Information 2018, 9, 149.

(55) Caruana, R.; Niculescu-Mizil, A. An empirical comparison of supervised learning algorithms. Proceedings of the 23rd international conference on Machine learning. 2006; pp 161–168.

(56) Sarkar, I. N.; Planet, P. J.; Desalle, R. caos software for use in character-based DNA barcoding. Molecular Ecology Resources 2008, 8, 1256–1259.

(57) Sievers, F.; Wilm, A.; Dineen, D.; Gibson, T. J.; Karplus, K.; Li, W.; Lopez, R.; McWilliam, H.; Remmert, M.; Söding, J.; Thompson, J. D.; Higgins, D. G. Fast, scalable generation of high-quality protein multiple sequence alignments using Clustal Omega. Molecular Systems Biology 2011, 7, 539.

(58) Swofford, D. Phylogenetic Analysis Using Parsimony. 2003,

(59) Virtanen, P. et al. SciPy 1.0: Fundamental Algorithms for Scientific Computing in Python. Nature Methods 2020, 17, 261–272.

(60) Li, D.-M.; Sun, H. TEP1, encoded by a candidate tumor suppressor locus, is a novel protein tyrosine phosphatase regulated by transforming growth factor *β*. Cancer research 1997, 57, 2124–2129.

(61) Liaw, D.; Marsh, D. J.; Li, J.; Dahia, P. L.; Wang, S. I.; Zheng, Z.; Bose, S.; Call, K. M.; Tsou, H. C.; Peacoke, M., et al. Germline mutations of the PTEN gene in Cowden disease, an inherited breast and thyroid cancer syndrome. Nature genetics 1997, 16, 64–67.

(62) Li, J.; Yen, C.; Liaw, D.; Podsypanina, K.; Bose, S.; Wang, S. I.; Puc, J.; Miliaresis, C.; Rodgers, L.; McCombie, R., et al. PTEN, a putative protein tyrosine phosphatase gene mutated in human brain, breast, and prostate cancer. science 1997, 275, 1943–1947.

(63) Song, Y.; DiMaio, F.; Wang, R. Y.-R.; Kim, D.; Miles, C.; Brunette, T.; Thompson, J.; Baker, D. High-resolution comparative modeling with RosettaCM. Structure 2013, 21, 1735–1742.

(64) Consortium, T. U. UniProt: a worldwide hub of protein knowledge. Nucleic Acids Research 2018, 47, D506–D515.

(65) Sunyaev, S. R.; Eisenhaber, F.; Rodchenkov, I. V.; Eisenhaber, B.; Tumanyan, V. G.; Kuznetsov, E. N. PSIC: profile extraction from sequence alignments with position-specific counts of independent observations. Protein engineering 1999, 12, 387–394.

(66) Denning, G.; Jean-Joseph, B.; Prince, C.; Durden, D. L.; Vogt, P. K. A short N-terminal sequence of PTEN controls cytoplasmic localization and is required for suppression of cell growth. Oncogene 2007, 26, 3930–3940.

(67) Nguyen, H.-N.; Yang, J.-M., Jr; Rahdar, M.; Keniry, M.; Swaney, K. F.; Parsons, R.; Park, B. H.; Sesaki, H.; Devreotes, P. N.; Iijima, M. A new class of cancer-associated PTEN mutations defined by membrane translocation defects. Oncogene 2015, 34, 3737–3743.

(68) Vousden, K. H.; Lu, X. Live or let die: the cell’s response to p53. Nature Reviews Cancer 2002, 2, 594–604.

(69) Cho, Y.; Gorina, S.; Jeffrey, P. D.; Pavletich, N. P. Crystal structure of a p53 tumor suppressor-DNA complex: understanding tumorigenic mutations. Science 1994, 265, 346–355.

(70) Olivier, M.; Hollstein, M.; Hainaut, P. TP53 mutations in human cancers: origins, consequences, and clinical use. Cold Spring Harb. Perspect. Biol. 2010, 2, a001008.

(71) Bromberg, Y.; Yachdav, G.; Rost, B. SNAP predicts effect of mutations on protein function. Bioinformatics 2008, 24, 2397–2398.

(72) Bromberg, Y.; Rost, B. SNAP: predict effect of non-synonymous polymorphisms on function. Nucleic acids research 2007, 35, 3823–3835.

(73) Porter, C. T.; Bartlett, G. J.; Thornton, J. M. The Catalytic Site Atlas: a resource of catalytic sites and residues identified in enzymes using structural data. Nucleic acids research 2004, 32, D129–D133.

(74) Doruker, P.; Atilgan, A. R.; Bahar, I. Dynamics of proteins predicted by molecular dynamics simulations and analytical approaches: Application to *α*-amylase inhibitor. Proteins: Structure, Function, and Bioinformatics 2000, 40, 512–524.

(75) Doss, C. G. P.; Zayed, H. Comparative computational assessment of the pathogenicity of mutations in the Aspartoacylase enzyme. Metabolic Brain Disease 2017, 32, 2105–2118.

(76) Studer, R. A.; Dessailly, B. H.; Orengo, C. A. Residue mutations and their impact on protein structure and function: detecting beneficial and pathogenic changes. Biochemical journal 2013, 449, 581–594.

(77) Mishra, S.; Singh, S.; Misra, K. Restraining pathogenicity in Candida albicans by taxifolin as an inhibitor of Ras1-pka pathway. Mycopathologia 2017, 182, 953–965.

(78) Senior, A. W. et al. Improved protein structure prediction using potentials from deep learning. Nature 2020, 577, 706–710.

